# Rapid evolution of flower phenology and clonality in restored populations of multiple grassland species

**DOI:** 10.1101/2022.10.28.514191

**Authors:** Anna Bucharova, Malte Conrady, Theresa Klein-Raufhake, Franziska Schultz, Norbert Hölzel

**Affiliations:** Department of Biology, Philipps University Marburg, Marburg, Germany; Institute of Landscape Ecology, University of Münster, Münster, Germany

**Keywords:** rapid evolution, ecosystem restoration, flowering phenology, seed-based restoration, adaptation

## Abstract

Restoration of terrestrial ecosystems often requires re-introduction of plants. In restored sites, the plants often face environment that differs from the one in natural populations. This which can affect plant traits, reduce performance and impose novel selection pressures. As a response, restored populations might rapidly evolve and adapt to the novel conditions. This may enhance population survival and contribute to restoration success, but has been rarely tested so far. Here, we focused on populations of three grassland species restored 20 years ago (*Galium wirtgenii, Inula salicina* and *Centaurea jacea*) by the transfer of green hay, and compared them with populations that were source of the hay. We measured plants both in-situ, and in common garden under control and three stress conditions. *In-situ*, restored and natural populations differed in flowering phenology in two out of the three species. In the common garden, plants of the restored population flowered earlier (in *Galium*) or showed increased plasticity of clonal propagation in response to clipping (*Inula*). Both these traits suggest rapid adaptation to the contrasting mowing regimes in restored in comparison to the natural donor sites. In *Centaurea*, we detected no differentiation, neither in-situ, nor in the common garden. Rapid evolution in two out of three species indicates that evolution in restoration may be rather common, yet not ubiquitous across species.

## Introduction

Restoration of degraded habitats is indispensable to combat current biodiversity crisis (Díaz et al., 2019). In terrestrial ecosystems, restoration often includes active planting or seeding to recover vegetation (Palma & Laurance, 2015). However, the environmental conditions at the degraded sites are commonly less favorable than the conditions in natural ecosystems (Brudvig et al., 2013), and introduced plants face novel challenges. This is especially true for habitat specialists, which are often the target of restoration efforts (Prach et al., 2013).

Environmental differences between natural and restored sites might be mirrored by intraspecific differences in plant phenotypes. For example, Klein-Raufhake et al. (2022) have shown that plants from grasslands restored 20 years ago are smaller and have different leaf chemical composition in comparison to the same species growing in natural populations, which correlates with the differences in soil chemistry due to post-agricultural legacy in restored sites. This pattern may be a simple plastic reaction to environmental conditions (Villellas et al., 2021). Alternatively, phenotypic differences can have a genetic basis – either because the introduced plant material genetically differed from plants in natural habitats (e.g. Aavik et al., 2012; Höfner et al., 2021; Kaulfuß & Reisch, 2019), or as a result of rapid evolution *in-situ*.

Plant populations can rapidly evolve if they face novel selection pressure and, at the same time, harbor heritable variability on which the new selection pressure can act (Etterson & Shaw, 2001). For example, *Brassica rapa* rapidly evolved towards earlier flowering in response to four years of drought (Franks et al., 2007), and similar patterns of drought adaptation have been documented over the past two decades in several other annual species (Rauschkolb et al., 2022). In the context of ecological restoration, first studies suggest that selection pressures at restoration sites can alter heritable phenotypic distribution of seeded species. Specifically, Kulpa & Leger (2013) detected strong selection towards smaller plant and seed size and earlier flowering in an annual grass, and Magnoli (2020) and Magnoli & Lau (2020) showed partly adaptive trait differentiation in an annual legume after seeding the same batch of seed to two restored sites. Yet, these studies focused on a single annual species each. The degree of evolution for the persistence of perennial plants in restored sites is unclear. Studies carried out in the framework of biodiversity experiments suggest that also perennial species evolve over less than two decades in response to community diversity (van Moorsel, Hahl, et al., 2018; van Moorsel, Schmid, et al., 2018). However, these experiments were controlled, with defined community composition and largely free from stochastic events. Whether the adaptive evolution of perennial species also happens in real-world restoration settings remains to be tested.

Here we focused on plants growing at restored and old seminatural meadows of the Upper Rhine floodplain, Germany. The old seminatural meadows have been initiated by humans centuries ago, and have been traditionally mown relatively late in the season for hay and stable litter. Despite these habitats have been created by humans and are dependent on human management, the long history of land use allowed the assembly of species-rich plant communities (Molinion, Cnidion). They are home of many specialized species and have a high conservation value (Bucharova et al., 2020). For simplicity, we call them “natural meadows” in in the following text. The natural meadows have been used as sources of diaspores for restoration of ca 70 ha of meadows at former arable land by the transfer of green hay. This approach is an established restoration method of central European meadows that transfers not only a large proportion of the target species, but also within-species genetic diversity of the donor populations (Dittberner et al., 2019; Hölzel & Otte, 2003; Kiehl et al., 2010).

Despite restoration efforts, restored and natural meadows differ in ecosystem functions and species composition. The soils of restored meadows contain a higher proportion of plant-available phosphorus and potassium (Donath et al., 2007; Hölzel & Otte, 2003). Consequently, they may be more productive and dominated by grasses, which implies higher competition for herbs (Figure S1). The management also differs: Natural meadows are mown mostly later in the season to mimic traditional management, and because the hay is used for ongoing restoration projects. Restored meadows are mown earlier, typically in mid June, to preserve nutritional value of the hay which is used as horse fodder. Apart from that these factors affect species composition (Busch et al., 2019; Poptcheva et al., 2009; Sehrt et al., 2020), they are also well documented selection pressures that drive plant local adaptation (Johnson et al., 2010; Rowe & Leger, 2011; Völler et al., 2013, 2017). It is thus possible that plants at restored sites evolved in response to the local conditions and diverged from the conspecifics growing at the source natural meadows. Such rapid evolution would allow adaptation to the less favorable environment of the restored sites and possibly contribute to the long-term restoration success.

We focused on three species that grow both at natural and restored meadows of the Upper Rhine floodplain, Germany: *Centaurea jacea, Galium wirtgenii* and *Inula salicina*. To evaluate the *in-situ* intraspecific variation between restored and natural sites, we scored phenology and performance traits *in-situ* in nearly 70 sites. To evaluate whether some of the differences are heritable, we collected seeds of each species from one natural site and a corresponding restored site that was established using hay sourced from the respective natural site, and grew them in a common garden site-by site. We also exposed the plants to nutrient limitation, competition and removal of aboveground biomass (to mimic mowing) to both simulate natural stressors and estimate possible adaptation. We hypothesize that 1) *in-situ*, plants growing in restored meadows differ from conspecific at natural sites, 2) part of this differentiation is heritable and 3) adaptive to environmental factors specific to natural and restored meadows.

## Methods

### Study sites

We focused on 17 natural meadows that served as hay donors for restoration, and 49 sites restored between 2000 and 2005. The restored sites were defined as the exact area where the hay has been transferred, and varied in size from ca 300m^2^ to ca 3000 m^2^. For each site, local authorities documented the year of restoration, as well as the natural site from which the hay was transferred. The restored sites are scattered in landscape of ca 3.5 km x 1km. As two adjacent restored sites are in some cases as close as 20 m, it is likely that there is gene flow between them.

### Study species

We focused on three species that grow both at natural and restored sites: *Galium wirtgenii, Centaurea jacea*, and *Inula salicina*, further referred by genus name. *Galium* is a short-lived perennial with limited clonal propagation, flowering in May and June. In contrast, *Inula* is a highly clonal perennial flowering only in July and August. *Centaurea* is a perennial species that does not spread clonally, and has a long flowering period from June to September. While *Galium* was present at all sites, the other two species were rarer: *Centaurea* was at 14 and 17, and *Inula* at 15 and 15 natural and restored sites, respectively.

### Field survey

We carried out the field survey over six days at the turn of May and June 2020. At each of the 49 restored and 17 natural sites, we randomly selected seven individual shoots of each target species per site. We kept at least 5 m between the shoots to avoid sampling close relatives or obvious clones. For each shoot, we measured height and phenology status. *Galium* was in the phase of early flowering at the time of sampling, with only part of the flowers being already fully open. We therefore estimated phenology as the proportion of fully open flowers, with more open flowers indicating earlier flowering. *Inula* had started to develop flower buds at the time of sampling. Thus, we estimated the phenology in three categories: (1) no flower bud yet, (2) developing flower bud, detected as a small ball hidden between the upmost leaves, and (3) developed flower bud of at least 4 mm size and with visible bracts. Although partly subjective, this approach was the only applicable to rate the apparent phenology differences between individuals at the time of sampling. *Centaurea* plants were all sterile at the time of sampling, and thus we did not record phenology.

### Seed collection

We collected seeds of the focal species for a common garden experiment in September 2019, shortly before the vegetation of natural sites was mown for green hay transfer for new restoration projects. As large parts of the restored sites were mown in the time of seed collection, we could effectively collect seeds only in stripes of vegetation that were left unmown as insect refugia, or, to lesser extent, plants that regrew after mowing. This limited the seed collection to one pair of restored and the corresponding natural site for each species, with the natural sites being exactly the ones from which the hay was originally sourced for restoration of the specific site. At each site, we collected seeds of 18-21 individuals per species, except *Centaurea* at the restored site where we could collect seeds from ten plants only.

### Common garden experiment

In March 2020, we sowed the seeds on seeding trays filled with seeding substrate located in a cold greenhouse (*Galium, Centaurea*). *Inula* seeds require a higher temperature for germination, therefore we placed the seeded trays to a 25°C warm climate cabinet and after germination to a cold greenhouse. When the seedlings developed the first true leaves, we transplanted up to 12 seedlings per field mother plant, depending on the germination rate, to quickpots filled with standard potting substrate. At the beginning of June, we transplanted the seedlings to 2.5L pots (filled with soil according treatment, see below), placed them in a common garden in a fully randomized design, and watered them regularly.

We assigned the plants to four treatments: control, competition, nutrient limitation and clipping (see below). To each treatment, we assigned 1-2 plants per field mother plant and species with the aim to have optimally 15-20 individuals per restoration status (natural vs. restored) and treatment. In *Galium*, the number of germinated seeds was not sufficient to apply all four treatments. Thus, we restricted the experiment to control and competition in this species. The final experiment contained 84 *Galium* plants divided into control and competition treatment, 119 *Centaurea* plants in all four treatments and 164 *Inula* plants, also in all four treatments, in total 367 plants.

The treatments were designed to meet the challenges faced by plants in their natural environment. Restored sites were more dominated by grasses and the productivity was higher and so is competition (Figure S1). To simulate this, we planted the focal individuals together with two seedlings of the grass *Lolium perenne*. The restored sites are mown early in summer, whereas the natural ones are mown later in summer or even in autumn. To simulate this, we clipped all aboveground biomass when approximately half of the individuals were flowering. The provision of soil nutrients typically differs between restored and natural meadows (Klein-Raufhake et al., n.d.). To simulate this, we planted the plants in nutrient-poor soil that contained 100 mg/l nitrogen, 26.2 mg/l phosphor and 91.3 mg/l potassium. We acknowledge that the low-nutrient treatment did not represent extremely poor soil compared to natural environments, but it was substantially poorer compared to standard soil used in the control and all other treatments (210 mg/l N, 109.5 mg/l P, 240.7 mg/l K). Control plants were grown alone, in standard in potting soil, and have not been clipped.

Throughout the season, we recorded flowering phenology three to four times a week as the day of the first open flower for each plant, starting with transplant date. We harvested the experiment at the end of August, when most of the control plants started to senescence. We measured the maximal height of each plant and counted the number of inflorescences (further called flowers for simplicity). In *Inula*, we recorded whether or not each plant created clonal runners. Finally, we harvested aboveground biomass, dried it at 70°C for 48 hours and weighted it.

### Statistical analyses

First, we analyzed field data to test for *in-situ* differences between plants at restored and natural sites. For each of the three species, we related height and flowering phenology (the latter except *Centaurea*) to population history (natural vs. restored) as fixed factor and to the site identity as a random factor to account for non-independency of plants growing at one site. The plant height in *Centaurea* and flowering phenology in *Inula* was log-transformed to approximate normal error distribution. We used linear mixed models (from the R-package lme4, Bates et al., 2015) with normal error distribution, and we graphically evaluated whether the model assumptions were met.

Second, we used the common garden data to test for heritable differences between plants from natural and restored meadows. The response variables were height, day of first flowering, number of flowers, and biomass in all three species, and presence of clonal runners in *Inula* only. We related respective response variable per species to population history (restored vs. natural) as fixed factor and maternal family as random factor to account for non-independency of plants from the same maternal family. As the variances within the treatments and population history were not homogenous, we estimated variance separately for each level of the respective factor using the function varComb of the R-package nlme (Pinheiro et al., 2018). Some response variables were log-transformed to meet the model assumptions (Table S2). For all response variables, we then assumed normal error distribution, except presence of clonal runners in *Inula*, where we used mixed models with binomial error distribution with the same random and fixed factors as above, but without accounting for non-equal variances. In all models, we graphically evaluated whether the assumptions were met.

To aid interpretation of the results, we tested for trait correlations of plants growing in the common garden. Specifically, we calculated a Pearson correlation matrix, between height, biomass, number of flowers and day of flowering in all species across all treatments except clipping. Plant regenerating after clipping were not included in the correlation because the treatment obviously reduced all trait values, which would confound the results. Clipping boosted clonal propagation in *Inula*, and significant interactions (see the results section for details) suggested trade-offs between clonality and height and biomass of the regenerating plants (Figure 3). We thus tested whether biomass and height of *Inula* that regenerated after clipping differed between plants with and without clonal propagation. We used Wilcoxon pair test to do this, as the assumptions for linear models were not met.

All analyses were done in R (R Development Core Team, 2020).

## Results

*In-situ*, both *Galium* and *Inula* showed more advanced flowering phenology in restored sites than in the natural ones (Figure 1, Table S1). Specifically, *Galium* had a higher proportion of open flowers and *Inula* showed a more advanced stage of flower bud development. To remind the reader, we did not assess flowering phenology in *Centaurea in-situ*, because all plants were still sterile at the time of field sampling. Plant height did not differ between natural and restored sites in any of the three focal species (Figure 1, Table S1).

**Figure 1:**
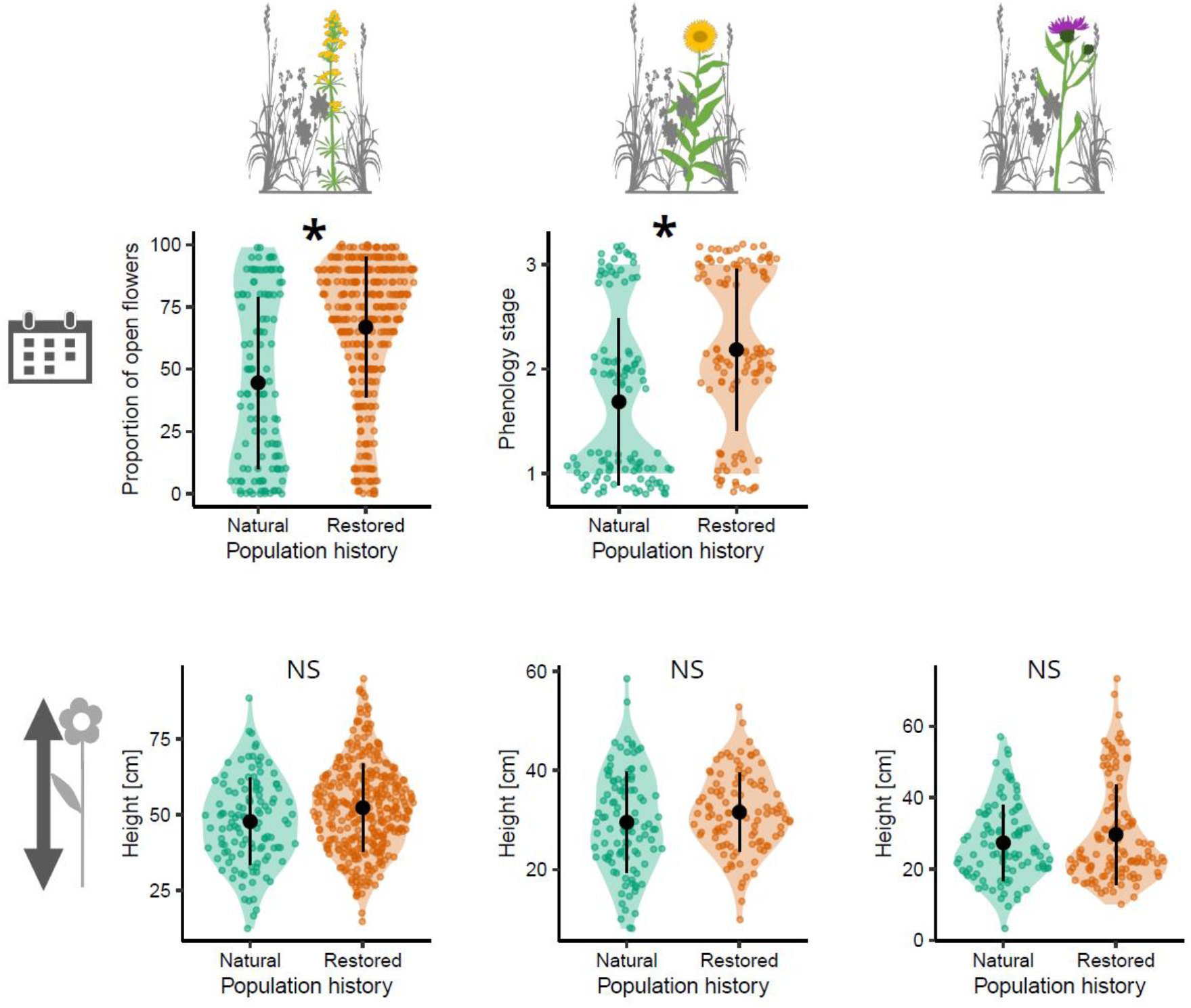
Data from the field. The difference in phenology status and height between plants growing at natural and restored sites, for *Galium* (left) and *Inula* (middle) and *Centaurea* (right). Black dots with whiskers represent mean ± SD, color dots measurements at individual plants. The star indicates significant differences (P<0.05), see Table S1 for detailed model results.

In the common garden, the stress treatments explained significant proportion of variability in biomass production and the number of flowers in all three species, and height in *Inula* and *Centaurea*. The stress treatments had no effect on plant flowering phenology (Figure 2, 3, S1, Table S2). Note that these are results of analysis of variance for the whole “treatment” variable, including all factor levels: control, competition, clipping and nutrient limitation (only the first two in *Galium*). We did not test specifically for effect of each stress treatment because it was not the focus of this study, and it is rather trivial that plants e.g. produce less biomass when they have less available nutrients. In *Galium*, plants from the restored site flowered earlier and produced more flowers than plants from the corresponding natural site. Significant interaction between population history and treatment in plant height indicated that plants from the restored and natural sites react differently to competition, specifically, plants from the natural site grew taller than the ones from restored site under control conditions but not under competition (Figure 2, Table S2). In *Inula*, none of the traits significantly depended on the population history as a main effect, but significant treatment : population history interaction in models for height, biomass and number of runners indicated that plants from restored and natural sites reacted differently to the stress treatments. Specifically, while plants from restored sites tended to be taller and produced more biomass under control treatment, the relation flipped when the aboveground biomass was previously clipped (Figure 3, Table S2). There was also a significant interaction in clonal reproduction (presence of clonal runners): Under control conditions, around 25% of plants produced clonal runners, independently of population history. The clipping treatment enhanced clonal propagation in all *Inula* plants, yet this increase was stronger in plants from restored (90% plant with runners) than from natural sites (63%), Figure 3, Table S2. In *Centaurea*, treatments explained significant proportion of variability in all traits except phenology, but neither the main effect of population history nor its interaction with treatments were significant (Figure S1, Table S2).

**Figure 2:**
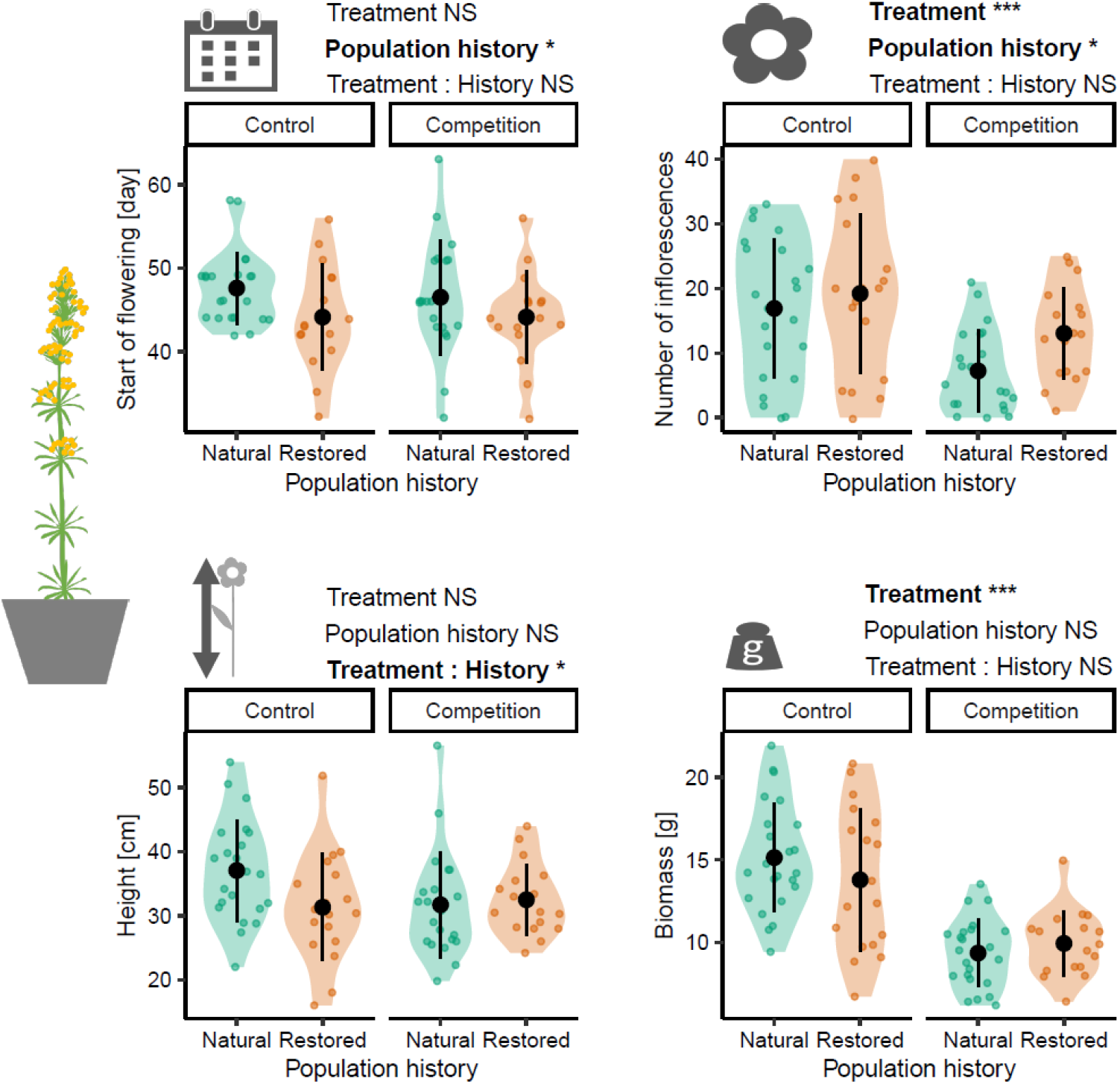
*Galium*, common garden. The difference between plants from restored and natural sites growing under control and competition treatment. Phenology (start of flowering), number of flowers, height and biomass. Short results of analysis of variance of the models including treatment (two factor levels), population history (two levels) and their interaction are above each plot, with significant terms in bold. Significance levels: NS non-significant, * *P* < 0.05, \*\**P* < 0.01, *** *P* <-0.001. See Table S2 for the full model results.

**Figure 3:**
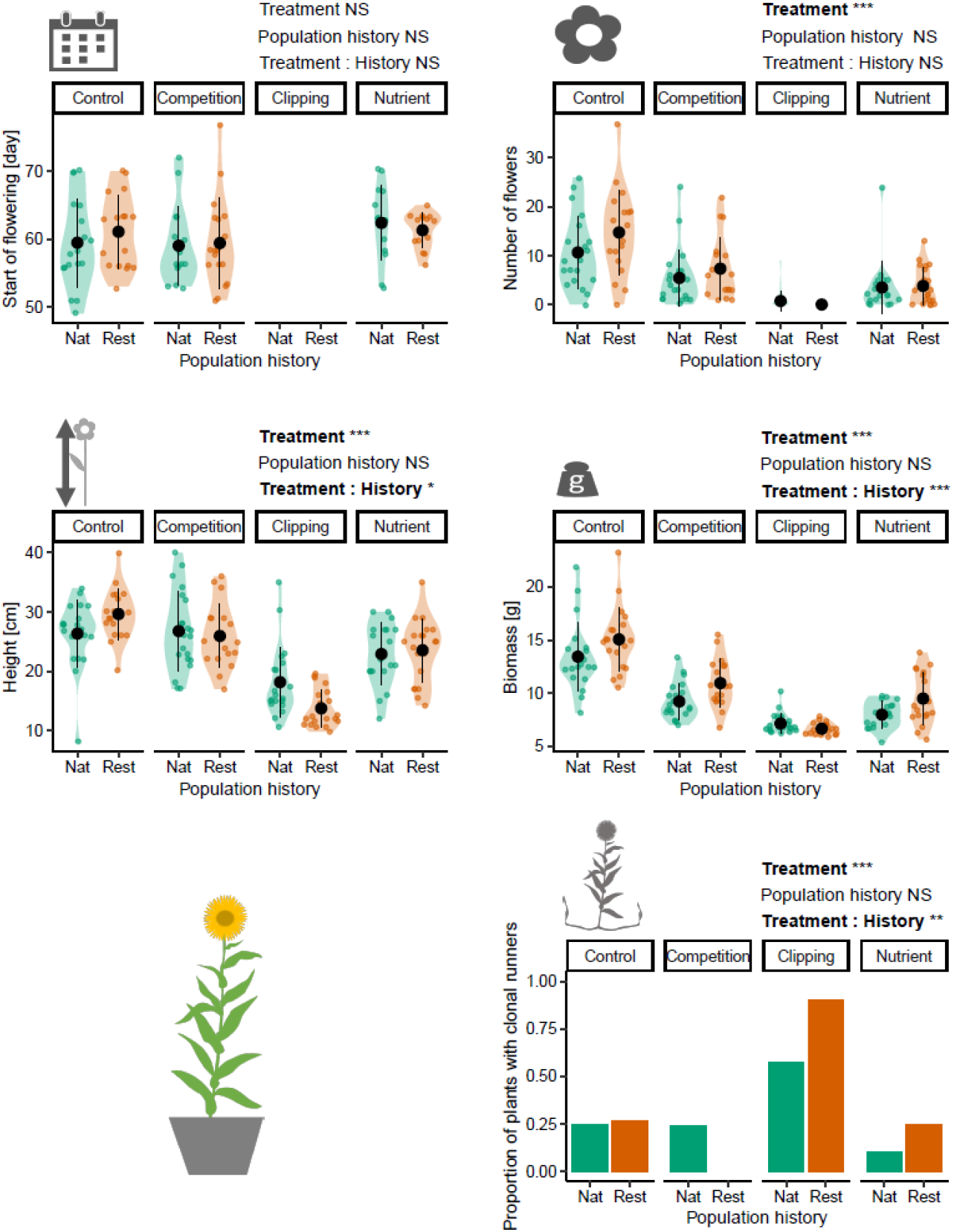
*Inula*, common garden. The difference between plants from restored and natural sites growing under the four treatments. Phenology (start of flowering), number of flowers, height, biomass and presence of clonal runners. Short results of analysis of variance of the models including explanatory variables treatment (four factor levels), restoration status (two levels) and their interaction are above each plot, with significant terms in bold. Significance levels: NS non-significant, * *P* < 0.05, \*\**P* < 0.01, *** *P* < 0.001. See Table S2 for the full model results.

Size-related traits, specifically biomass, height and number of flowers were positively correlated with each other in all three species growing in the common garden (Figure 4, Figure S2). These size-related traits were negatively correlated with the start of flowering in *Galium* and *Inula*, which means that the larger plants flowered earlier. There was no such correlation in *Centaurea* (Figure S2). *Inula* plants regenerating after clipping were smaller (P<0.001) and had less aboveground biomass (P=0.024) in case they produced clonal runners.

**Figure 4:**
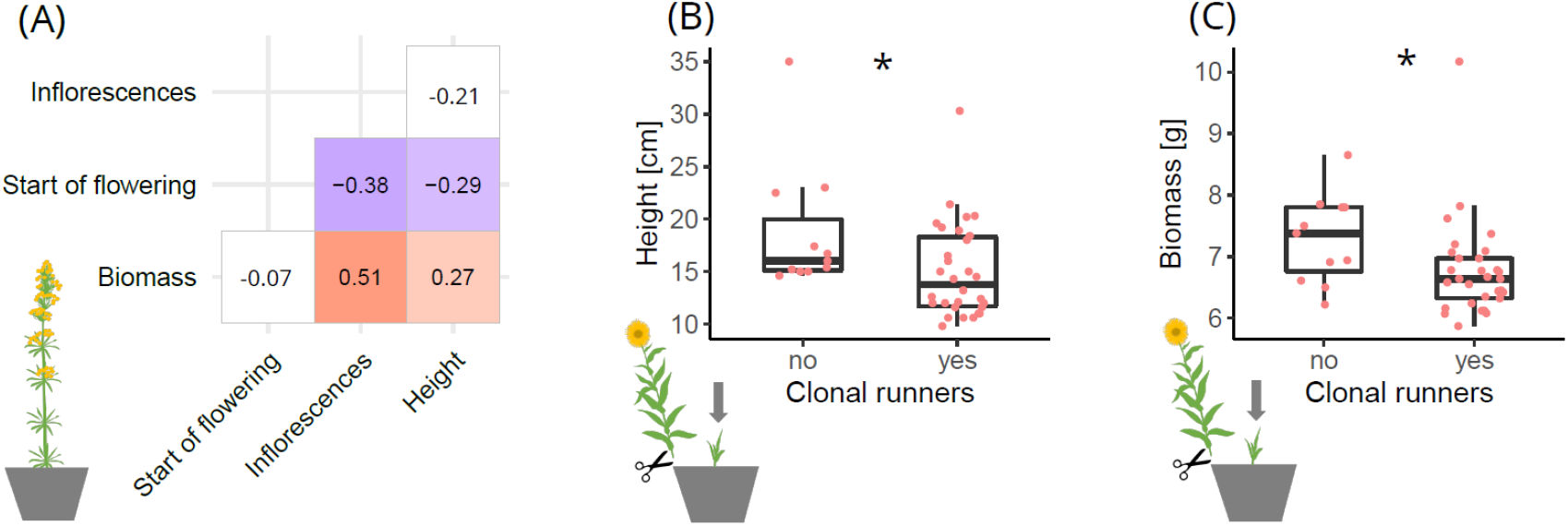
(A) Correlation among traits in *Galium* in common garden. Numbers indicate correlation coefficients, significant values(P<0.05) are highlighted in color. Trait correlations for the other two species are in Figure S2. (B, C) *Inula*, common garden, plants regenerating after clipping, height and biomass in plants that did and did not produce clonal runners. Stars indicates significant difference (P<0.05).

## Discussion

Revegetation is an indispensable part of many terrestrial restoration projects. Yet, plants introduced to degraded habitats face challenging environments that can affect their traits and fitness. Such trait changes can be a simple plastic reaction to the novel conditions. Alternatively, novel selection pressures may push heritable variation of the restored population towards phenotypes that are better adapted to the restoration condition; a process that can contribute to restoration success. Here we show that phenotypes of *Galium wirtgenii* and *Inula salicina* growing *in-situ* in meadows restored 20 years ago differ from conspecifics in natural meadows that served as source for restoration. A common garden experiment revealed that these differences are partly heritable and likely adaptive, indicating adaptive evolution. Evolution in restoration was documented previously in two annual species (Kulpa & Leger, 2013; Magnoli, 2020). We bring further evidence for two perennials, suggesting that evolution in restoration is probably rather common. However, not universal, as in the third species we studied, *Centaurea jacea*, there was no differentiation between restored and natural populations.

*Galium wirtgenii* in restored meadows evolved towards earlier flowering. *In-situ, Galium* flowered earlier in restored meadows than conspecifics in natural sites. Common garden plants showed the same pattern, indicating that the difference in flowering phenology is heritable. Plants from restored populations also produced more flowers in the common garden, but as the number of flowers correlated with earlier flowering, these two traits are part of the same syndrome. Earlier flowering of restored populations is likely a result of adaptation to mowing regime. Natural meadows that were the source of hay for restoration are typically mown in late summer or even autumn, but restored meadows are mown in mid-June when *Galium* starts to set seeds. Only genotypes that produce ripe seeds before mowing are able to contribute to the next generation. Mowing thus creates effective selection pressure and shifts the restored population towards earlier flowering. In fact, flowering time is well known as a key trait in local adaptation to various mowing and grazing regimes (Reisch & Poschlod, 2009; Völler et al., 2013, 2017), yet we are the first to document this process in recently restored grasslands.

In contrast to *Galium*, we did not detect evolution of flowering time in *Inula salicina*. The species showed advanced phenology at restored sites *in-situ*, but this was a simple plastic reaction to environmental conditions, because there was no difference in the common garden. Restored meadows are less surrounded by trees and thus less shaded, which can cause a warmer microclimate and result in earlier flowering (Willems et al., 2021). The lack of an evolutionary response in flowering time can be explained by the life history traits of the species, specifically late flowering and strong clonal reproduction. In natural meadows, the species flowers in summer and sets seeds before mowing in autumn. Plants in restored sites are mostly mown in June, before the seed set, and generative reproduction is limited to the rare years when there is enough summer rain to allow regrowth and second flowering in autumn. Restored populations are thus likely maintained mainly by clonal propagation.

The restored population of *Inula salicina* evolved enhanced plasticity of clonal reproduction in response to mowing. Clipping in the common garden boosted the clonal reproduction in this species, a reaction that is common in grassland clonal plants (Tachibana et al., 2010). Yet, this response was much stronger in plants from restored sites (Figure 3), likely as an adaptation to early mowing which limits generative reproduction there. Plants from restored sites were also taller and produced more biomass, but this relationship flipped when the plants regenerated after clipping. This is likely a result of a trade-off between aboveground size and clonal runners, as plants grew shorter when they invested into clonal propagation after clipping (Figure 5). Clonal propagation in general is a common adaptation of species that grow in frequently mown or grazed habitats (Wellstein & Kuss, 2011), yet within-species adaptive differentiation in ability of clonal reproduction was so-far only rarely documented (e.g. Simon-Porcar et al., 2021; Wang et al., 2016).

In the third species, *Centaurea jacea*, we did not detect any differentiation, neither in the field nor in the common garden. The species flowers only after the restored meadows are mown, and there is thus no opportunity for selection. It also grows in a wide range of conditions (Leuschner & Ellenberg, 2018), and is one of the few species that can thrive in grassland restorations in unfavorable postarable soils (Klein-Raufhake et al., 2022.). Thus, it is possible that *Centaurea* has such a wide ecological niche that the environment at restored meadows is not challenging. Substantial reaction to stress treatments in common garden also suggests that many traits of the species are rather plastic, which might help to maintain fitness in both restored and natural habitats.

The evolution in restoration could be affected by gene-flow from surrounding populations. The baseline for each restored population was the genetic composition of the natural population that was the source for the hay transfer (Dittberner et al., 2019). Heritable differentiation between natural and restored populations could rise by sole selection on the extant genetic variability, as it was the case in a previous study focusing on evolution in restoration (Magnoli, 2020). However, in the present system, other conspecific populations are nearby and post-restoration gene flow is likely (Höfner et al., 2021). This is especially true in *Galium*, which relies on generative reproduction and gene flow could be via both seed and pollen. The contribution of gene flow to the observed phenotypic differentiation between populations is hard to predict. On one hand, gene flow is largely thought to counteract adaptive evolution (Sexton et al., 2014). On the other hand, gene flow might have increased genetic variability on which selection can act (Barrett & Schluter, 2008) and thus facilitate adaptation. Understanding the contribution of post-restoration gene flow among populations in this system will require further studies, optimally using genomic methods.

Our findings indicate that plants can adapt to novel conditions in restoration, yet certain caution is necessary when interpreting the results. First, we worked with seeds collected directly from the field, which could be affected by maternal effects. Maternal effects influence mainly early life stages such as germination and seedling establishment. Adult traits, as measured in this study are less likely to be affected (Weiner et al., 1997). Second, we studied only one pair of restored and natural population for each species, and the results could be population-specific. To achieve more general findings, future studies should include multiple populations. However, this will come at the cost of studying multiple species, because all experiments are limited in size.

### Implication for practice

Active revegetation is indispensable for numerous terrestrial restoration projects. Here we show that even when the plant diaspores come from local natural populations, the traits of plants in restored populations differ from the ones growing in the source natural populations. Part of this differentiation is plastic reaction to the environment, but in two out of three species we detected also heritable and putatively adaptive changes. This suggests that the restored populations are not static. There is a substantial evolutionary dynamic that has the potential to improve adaptation of introduced material to the on-site condition, and possibly enhance restoration success. Such eco-evolutionary dynamics, however, can act only in presence of genetic diversity. Here, enough genetic variability was provided in material sourced from local natural populations to allow adaptation to novel management. However, in the case of restoration projects in highly degraded habitats or in regions with predicted strong environmental change, it might be a good idea to implement seed sourcing strategies that deliberately enhance genetic diversity of seed, either of wild collected or agriculturally propagated material (Bucharova et al., 2019; Conrady et al., 2022).

## Aknowledgement

We thank M. Harnisch, Administration of Riedstadt town, for information on restoration project, V. Ratka, J. Klose, L. Nährtke, T. and T. Saracoglu for technical assistance.

## Author contribution

AB conceived the idea; AB and NH designed the study; FS collected the field data; FS, TKR and MC performed the experiment; AB analyzed the data and wrote the first draft; all authors contributed to editing of the final text.

## Supplementary material

**Table S1:** Results of the models testing the difference in phenology and height of the three focal species growing in natural vs restored meadows. significant values are in bold.

|  | Phenology |  |  | Height |  |  |
| --- | --- | --- | --- | --- | --- | --- |
|  | df | F | P | df | F | P |
| *Galium* | 1, 63.23 | 28.15 | **<0.001** | 1, 64.32 | 3.3 | 0.073 |
| *Inula* | 1, 28.01 | 10.54 | **0.003** | 1, 28.08 | 0.67 | 0.42 |
| *Centaurea* | NA | NA | NA | 1, 29.00 | 0.34 | 0.554 |

**Table S2:** Results of the models testing the effect of stress treatments (control, drought, low nutrient, clipping) on plants from natural and restored sites (source habitat) in common garden on traits of the three focal species. Degrees freedom (df) are indicated in format nominator, denominator df. For testing phenology differences, clipping treatment was not included, since many plants have been clipped before they started flowering and the phenology variable is thus meaningless in this treatment. Significant values are in bold.

| *Galium* | Start of flowering |  |  | Nr. of flowers |  |  | Height (log) |  |  | Biomass |  |  |
| --- | --- | --- | --- | --- | --- | --- | --- | --- | --- | --- | --- | --- |
|  | df | F | P | df | F | P | df | F | P | df | F | P |
| Treatment | 1, 52 | 0.34 | 0.563 | 1, 55 | 17.01 | **<0.001** | 1, 55 | 1.26 | 0.266 | 1, 58 | 59.12 | **<0.001** |
| Source habitat | 1, 52 | 5.11 | **0.028** | 1, 55 | 6.31 | **0.014** | 1, 55 | 1.11 | 0.298 | 1, 58 | 0.04 | 0.848 |
| Treatment:Source | 1, 52 | 0.16 | 0.688 | 1, 55 | 0.73 | 0.395 | 1, 55 | 4.51 | **0.038** | 1, 58 | 1.97 | 0.166 |

| *Inula* | Start of flowering |  |  | Nr. of flowers (log) |  |  | Height |  |  | Biomass (log) |  |  |
| --- | --- | --- | --- | --- | --- | --- | --- | --- | --- | --- | --- | --- |
|  | df | F | P | df | F | P | df | F | P | df | F | P |
| Treatment | 2, 70 | 1.75 | 0.181 | 3, 130 | 120.65 | **<0.001** | 3, 130 | 62.63 | **<0.001** | 3, 129 | 136.3 | **<0.001** |
| Source habitat | 1, 70 | 0.00 | 0.990 | 1, 130 | 0.29 | 0.590 | 1, 130 | 0.89 | 0.348 | 1, 129 | 0.82 | 0.376 |
| Treatment:Source | 2, 71 | 0.58 | 0.565 | 3, 130 | 2.13 | 0.100 | 3, 130 | 4.65 | **0.004** | 3, 129 | 6.21 | **<0.001** |

| *Centaurea* | Start of flowering |  |  | Nr. of flowers (log) |  |  | Height |  |  | Biomass (log) |  |  |
| --- | --- | --- | --- | --- | --- | --- | --- | --- | --- | --- | --- | --- |
|  | df | F | P | df | F | P | df | F | P | df | F | P |
| Treatment | 2, 45 | 1.69 | 0.196 | 3, 89 | 125.40 | **<0.001** | 3, 89 | 103.24 | **<0.001** | 3, 89 | 429.54 | **<0.001** |
| Source habitat | 1, 45 | 0.31 | 0.578 | 1, 89 | 0.18 | 0.673 | 1, 89 | 0.34 | 0.563 | 1, 89 | 0.75 | 0.390 |
| Treatment:Source | 2, 45 | 0.45 | 0.643 | 3, 89 | 0.80 | 0.496 | 3, 89 | 0.22 | 0.883 | 3, 89 | 0.84 | 0.475 |

| *Inula* | Clonal runners (yes/no) |  |  |
| --- | --- | --- | --- |
|  | df | Chi-square | P |
| Treatment | 3 | 41.78 | **<0.001** |
| Status | 1 | 1.06 | 0.304 |
| Treatment:Source | 3 | 13.2 | **0.004** |

**Figure S1:**
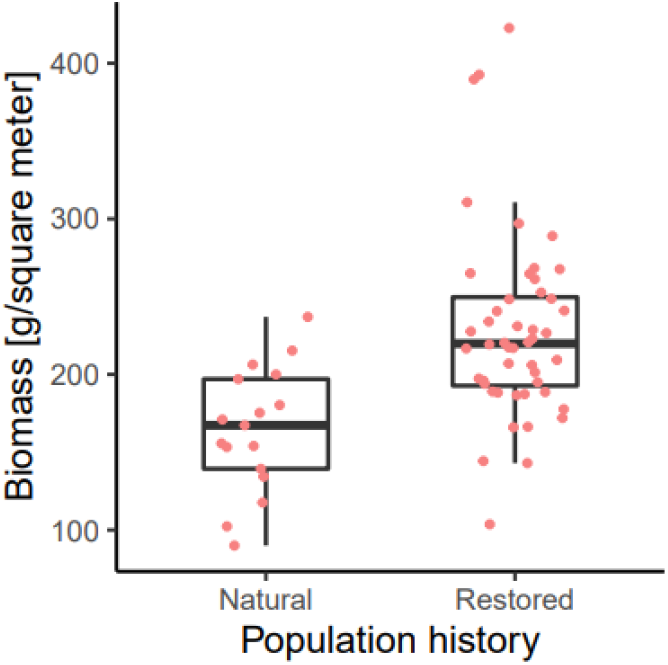
The difference in biomass productivity between natural and restored meadows. At each site, we randomly placed three 0.1m^2^ quadrant and cut all aboveground biomass, pooled it into one bag, dried it at 70°C and weighted. To test for the difference, we fit a linear mixed model with biomass as response variable and population history es explanatory variable. Restored meadows produced significantly more biomass than natural meadows (R^2^= 0.192, *P*<0.001).

**Figure S2:**
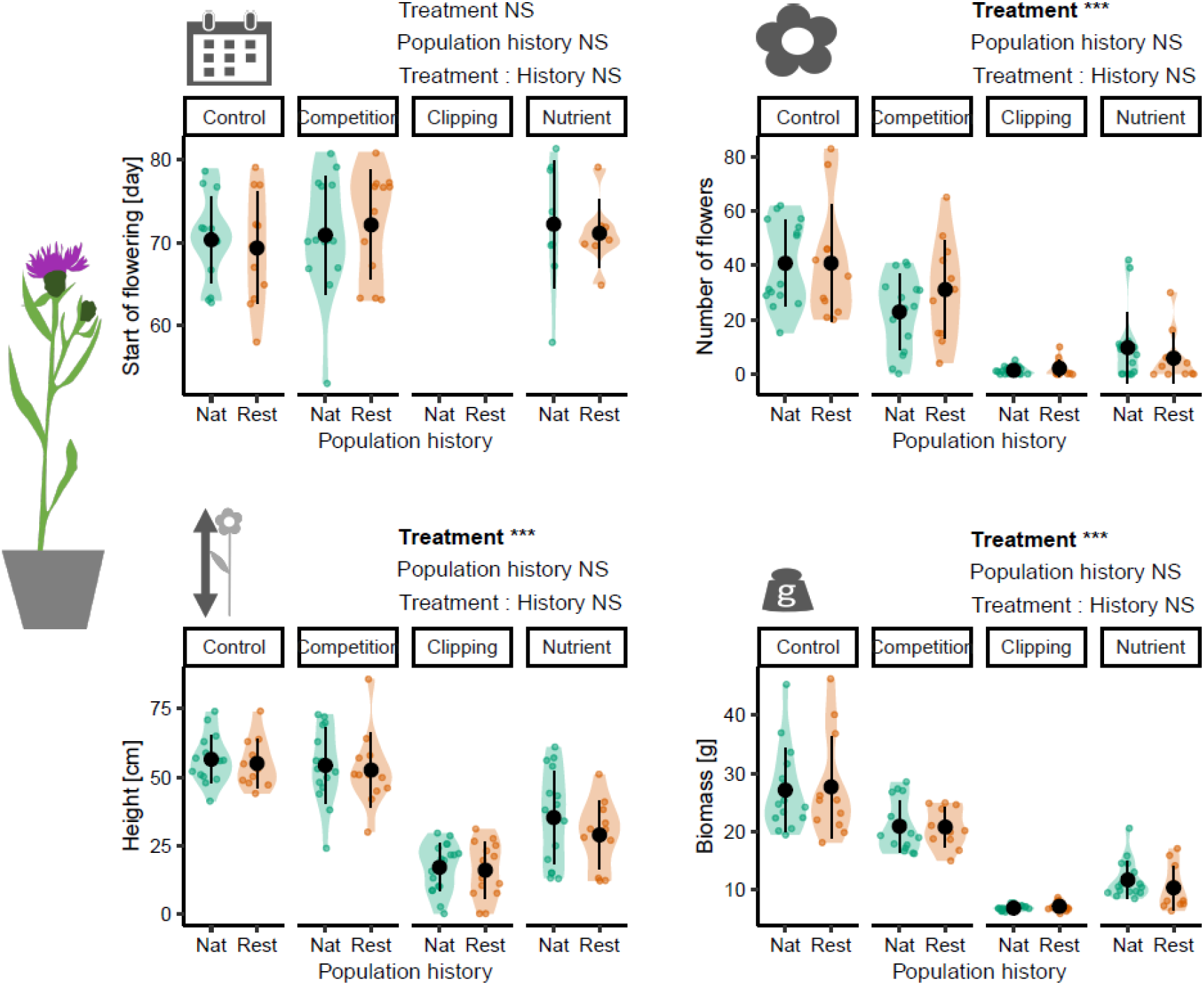
*Centaurea*, common garden. The difference between plants from restored and natural sites growing under the four treatments. Phenology (day of flowering), number of flowers, height and biomass. Short results of analysis of variance of the models including explanatory variables treatment (four factor levels), restoration status (two levels) and their interaction are above each plot, with significant terms in bold. Significance levels: NS non-significant, * *P* < 0.05, \*\**P* < 0.01, *** *P* <-0.001. See Table S3 for the full model results.

**Figure S3:**
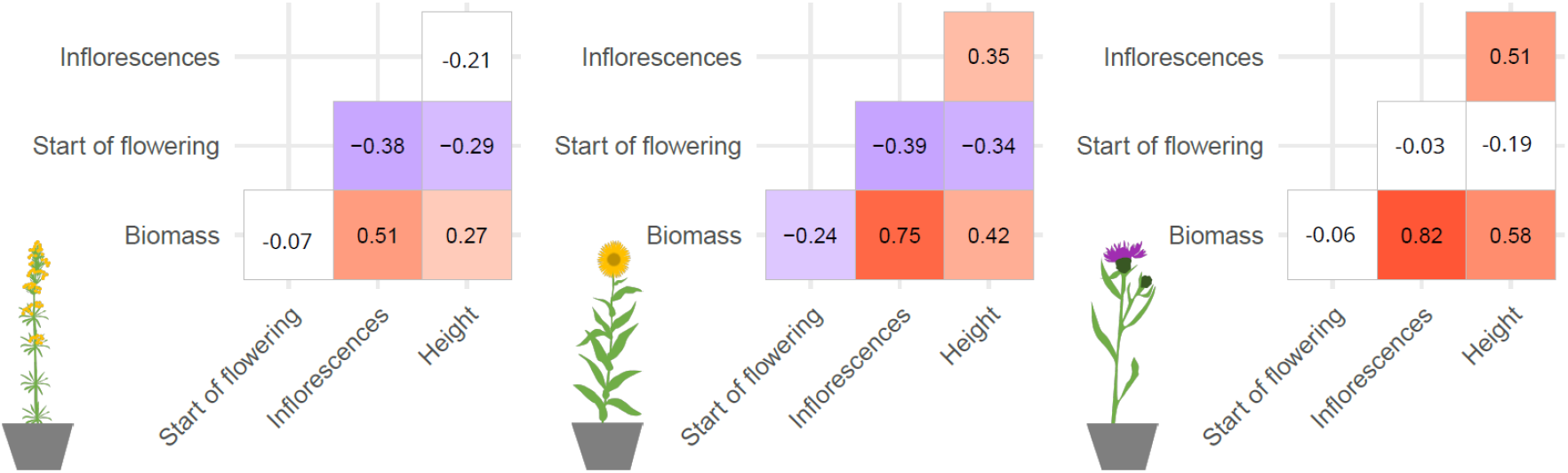
Correlation among traits in *Galium* (left), *Inula* (middle), and *Centaurea* (right) growing in the common garden. Numbers indicate correlation coefficients, significant values(P<0.05) are highlighted in color.

## Notes

### Competing Interest Statement

The authors have declared no competing interest.

